# Predicting Gene Mutations in Colon Cancer Using Long-Term Temporal Dependency Learning on a Directed Co-Occurrence Asymmetry Graph

**DOI:** 10.64898/2026.01.22.701121

**Authors:** Collin Sumrell, Aleksandr Shishkin, Ekene Okeke, Alexander Zelikovsky, Marmar R. Moussa

**Affiliations:** School of Computer Science, University of Oklahoma, Norman, OK 73019, United States; Department of Computer Science, Georgia State University, Atlanta, GA 30303, United States; Stephenson School of Biomedical Engineering, University of Oklahoma, Norman, OK 73019, United States

**Keywords:** somatic mutation, colorectal cancer, cancer genomics, directed acyclic graph, long short-term memory

## Abstract

**Motivation:** Colorectal tumorigenesis follows stepwise mutational progression patterns, but inferring dependencies from mutation data remains challenging. We propose a directed co-occurrence asymmetry graph to infer graph-derived mutational paths from 2,344 colon adenocarcinoma samples covering 23,858 mutated genes.

**Results:** We introduce a gene-conditioned shared Long Short-Term Memory (LSTM) model with attention to predict mutation status along inferred paths. The attention mechanism learns to weight informative predecessor mutations directly, while gene embeddings adapt the shared model to each target gene. We compare this model with standard per-gene LSTM and dilated Convolutional Neural Network (CNN) architectures. Graph-derived paths substantially improved precision and recall over the reproduced frequency-ordering baseline, with weighted paths performing best overall. The attention-based shared LSTM achieved competitive area under the ROC curve (AUC) and high recall, indicating that attention over predecessor mutations provides a useful representation for target-gene prediction.

**Code and data:** github.com/moussa-lab/MutationPrediction

## 1 Introduction

Colorectal adenocarcinoma is an evolutionary process driven by the stepwise accumulation of somatic mutations. While most somatic mutations are passengers that do not confer a selective advantage, a subset of driver mutations promotes tumorigenesis and progression (Vogelstein *et al*. 2013). This process has often been modeled as a temporal sequence of events. For example, the Fearon-Vogelstein model proposes that colorectal adenocarcinoma typically begins with inactivation of *APC*, followed by mutations in *KRAS* and *TP53* (Fearon and Vogelstein 1990). Understanding these temporal dependencies is important because early drivers can shape tumor evolution, and mutations along the disease progression pathway influence patient prognosis. However, while well-studied genes like *APC* display strong temporal signals, the relationships between less frequent mutations, both driver and passenger, remain poorly understood (Kumar *et al*. 2020).

Previous studies have inferred a global gene order using frequency-based approaches and then used these orders to train deep learning models of cancer evolution. This strategy was applied by Auslander *et al*. (2019) using Long Short-Term Memory (LSTM) networks to learn mutational load and perform next-step prediction. Using a pseudotemporal gene ranking based on mutational frequency ratios, the authors trained LSTMs to predict the mutational burden and the occurrence of future drivers with high accuracy. However, our attempt to reproduce the mutation-prediction component of Auslander *et al*. (2019) yielded recall and precision metrics below a random-guessing baseline. Because these mutation matrices are highly sparse, the prediction task is inherently class imbalanced: mutations are rare events, so frequently predicting no mutation can yield high accuracy while producing poor true positive rates. Another limitation of Auslander *et al*. (2019) was the reliance on a single global, frequency-based formula ordering of all genes detected in the study cohort. Furthermore, the study used a relatively limited number of colorectal cancer (CRC) samples for model training (*n* = 253).

Despite the utility of a global sequential gene order, a single mutation can drive multiple downstream mutations simultaneously, creating a complex network of mutational dependencies. Indeed, in CRC, tumors often develop from a series of genetic changes described by two major pathways: the ‘conventional adenoma-carcinoma sequence’, with mutations in *APC* considered a classic driver mutation (Jones *et al*. 2008); and a secondary, less known ‘alternative serrated pathway’, which in turn branches into various paths that are often driven by mutations in either *KRAS* or *BRAF* and leading to different consensus molecular subtypes (Fessler *et al*. 2016; IJspeert *et al*. 2017) of the same disease.

This network-like mutational complexity, together with the limitations of existing frequency-based orderings, motivated a graph-based approach for inferring data-driven mutational orders. Rather than modeling tumorigenesis as a fixed linear sequence, we represent mutational progression as a directed acyclic graph (DAG). In this graph, genes are represented as vertices and asymmetric co-occurrence relationships are represented as directed edges, allowing the model to capture multiple potential evolutionary pathways.

We further propose a prediction framework that combines graph pathway inference with attentive, gene-conditioned sequence modeling. We first construct a directed graph of gene relationships based on conditional probabilities observed in a large cohort of human colon cancer samples. We hypothesize that weighted and unweighted longest paths through this directed co-occurrence graph represent dominant potential evolutionary trajectories of the disease.

Using the inferred paths, we introduce a shared LSTM with attention and learned gene embeddings for downstream mutation prediction. Unlike independent per-gene models, this architecture pools training examples across target genes while conditioning each prediction on gene identity. Prior work has shown that attention can improve recurrent sequence classification by emphasizing informative input positions (Shen and Lee 2016); here, attention allows the model to weight informative predecessor mutations directly rather than relying only on the final LSTM state. We compare this proposed model with conventional per-gene LSTM and dilated Convolutional Neural Network (CNN) architectures to evaluate whether the graph-derived paths provide useful predictive context.

In our approach, each tumor is treated as a cross-sectional binary snapshot, and pseudotemporal ordering is inferred from inter-patient co-occurrence asymmetries. As a result, the inferred paths should be interpreted as population-level statistical ordering signals rather than patient-specific evolutionary timelines. We use sequence models as a predictive validation mechanism: for each target gene at position *t*, the model evaluates whether that gene is mutated from the mutation states of genes inferred to occur earlier in the trajectory. The LSTM and dilated CNN models are used to evaluate whether the graph-derived ordering provides useful predictive context, not for graph or pathway inference.

## 2 Materials and methods

### 2.1 Data preparation

Raw data from nine datasets were obtained through cBioPortal (Cerami *et al*. 2012; Gao *et al*. 2013; de Bruijn *et al*. 2023), consolidated into one large cohort, and binarized into a matrix **M** representing the somatic mutation status of each gene. For gene *i* in sample *k*, the entry *m*_*ik*_ was set to 1 if a mutation was detected and 0 otherwise. We retained only genes mutated at least once across the constituent datasets and dropped all others, resulting in 23,858 genes across 2,344 samples. The mutation distribution for the 20 most frequently mutated genes is shown in Figure 1. The constituent datasets of **M** were:

**Figure 1.**
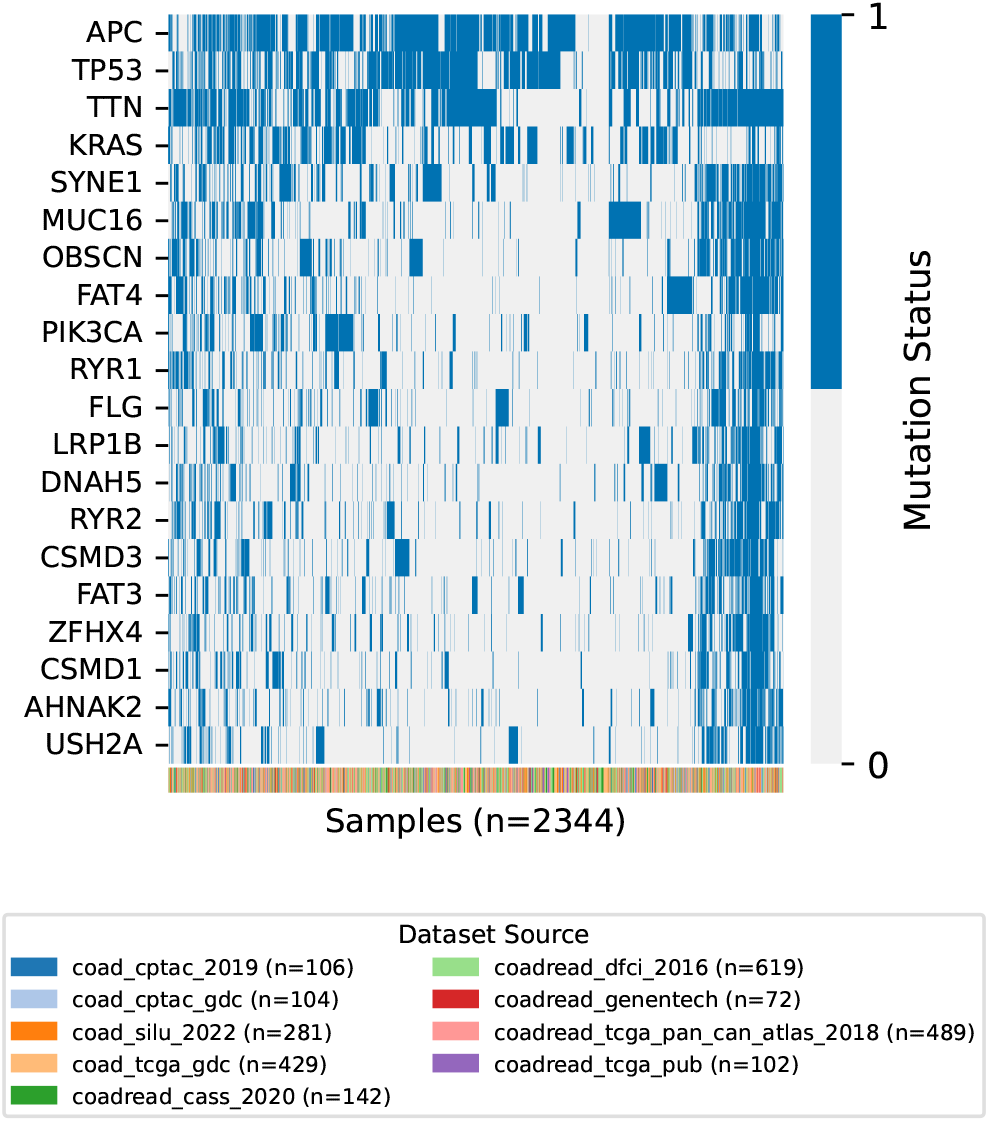
Binary mutation statuses (*m*_*ik*_) for the 20 most frequently mutated genes across 2,344 samples. Rows correspond to genes sorted by overall mutational prevalence, while columns represent patient samples ordered using hierarchical clustering (Euclidean distance, average linkage) to show patterns in mutational burden and co-occurrence. Blue indicates the presence of a somatic mutation; light gray indicates the absence of a mutation. The color-coded annotation bar at the bottom identifies the originating dataset for each sample.

- coad_cptac_2019 (*n* = 106 samples) (Vasaikar *et al*. 2019)
- coad_cptac_gdc (*n* = 104) (Heath *et al*. 2021)
- coad_silu_2022 (*n* = 281) (Roelands *et al*. 2023)
- coad_tcga_gdc (*n* = 429) (Heath *et al*. 2021)
- coadread_cass_2020 (*n* = 142) (Li *et al*. 2020)
- coadread_dfci_2016 (*n* = 619) (Giannakis *et al*. 2016)
- coadread_genentech (*n* = 72) (Seshagiri *et al*. 2012)
- coadread_tcga_pan_can_atlas_2018 (*n* = 489) (Heath *et al*. 2021)
- coad_tcga_pub (*n* = 102) (Cancer Genome Atlas Network 2012)

### 2.1.1 Clustering vs. clinical stage

Of the 2,344 samples, 1,516 included American Joint Committee on Cancer (AJCC) TNM staging information. TNM staging describes disease extent in terms of local primary tumor growth (T), regional lymph node involvement (N), and distant metastatic spread (M), with stages ranging from I for more localized disease to IV for the most advanced disease (Amin *et al*. 2017). To test whether somatic mutation profiles reflect clinical stage and whether stage information should be incorporated into downstream modeling, we performed unsupervised clustering on the staged samples and compared the resulting groupings to AJCC stages using the adjusted Rand index (ARI) (Hubert and Arabie 1985). The analysis was conducted on both the full feature set (23,858 genes) and a filtered subset of genes mutated in at least 5% of samples (1,453 genes). We then applied four clustering strategies: Jaccard distance hierarchical clustering with Ward linkage (Ward 1963) (*k* = 4), Louvain community detection (Blondel *et al*. 2008) on a KNN graph (*k* = 15) constructed in PCA-reduced space, Leiden community detection (Traag *et al*. 2019) on the same graph, and Ward hierarchical clustering in PCA space with *k* chosen by silhouette score. The Pearson correlation between PC1 and tumor mutational burden was also computed. Additional exploratory analyses of filtered and unfiltered clustering results, as well as stage- and Louvain-stratified graph-derived prediction experiments, are provided in Supplementary Materials.

### 2.2 Graph construction

To model potential dependency relationships between mutations, we computed an asymmetric conditional probability metric for each pair of genes. We defined the probability of observing mutation *i* given mutation *j* as:

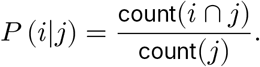

We directed an edge from *i* → *j*, interpreted as mutation *i* preceding mutation *j*, when *P* (*i*|*j*) > *P* (*j*|*i*). This condition means that samples with mutation *j* usually also contain mutation *i*, whereas mutation *i* is more often observed without mutation *j*; in other words, *j* tends to appear only after *i* has already occurred. Using the counts *c*_*ab*_, where *a* indicates the absence/presence of mutation *i*_*c* +*c*_ and *b* indicates the absence/presence of mutation *j*, these probabilities are 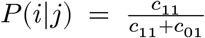, 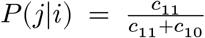Therefore, *P* (*i*|*j*) > *P* (*j*|*i*) corresponds to *c*_01_ < *c*_10_: mutation *j* rarely occurs without *i*, while mutation *i* more often occurs without *j*.Edge weights *w*_*ij*_ were assigned according to the magnitude of the conditional probability asymmetry:

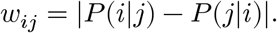

To reduce noise, potential edges with weight *w*_*ij*_ < ϵ were excluded from consideration (all reported results use ϵ = 0.02). To ensure consistency in ordering, we refined the graph into a directed acyclic graph (DAG) by removing the minimum set of edges required to break cycles, using the feedback arc set algorithm (Eades *et al*. 1993). This approach minimizes the total weight of removed edges, preserving the strongest inferred dependencies while eliminating logical contradictions in the pathway structure.

### 2.3 Bootstrap consensus graph construction

As an alternative to the primary graph construction approach, we also explored a bootstrap aggregation procedure to assess a consensus graph. In this approach, 200 bootstrap datasets were generated from the COAD training cohort used by Auslander *et al*. (2019), rather than from the full curated mutation matrix **M**. A directed graph was constructed from each of those datasets using the conditional probability and asymmetry framework described above, with edges below a probability threshold of 0.05 excluded.

After processing all bootstrap datasets, edges were aggregated into a consensus edge list recording each edge’s frequency *f*_*ij*_, defined as the number of bootstrap datasets in which the edge appeared, and its mean weight 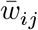 across those datasets. Rather than applying the feedback arc set algorithm to enforce acyclicity, we identified the minimum frequency threshold *T*_min_ such that retaining only the edges appearing in at least *T*_min_ bootstrap datasets produced a valid DAG.

### 2.4 Pathway inference

We hypothesized that dominant evolutionary trajectories would correspond to longest paths within a DAG. To that end, we computed two longest paths for each graph. The first, an unweighted longest path, maximized the number of vertices in the path. The second, a weighted longest path, was computed by identifying the sequence of vertices that maximized the total edge weight, 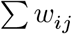 or total mean edge weight, 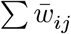, through the graph. We focused on these two paths because they capture distinct properties of the DAG. They do not exhaust the possible trajectories in the graph, but provide two interpretable candidates for evaluating whether graph-derived mutation orders support downstream prediction.

### 2.5 Prediction task formulation

We formulated mutation-status prediction as a sequence modeling task. For each target gene, indexed by its pseudotemporal position *t* within the inferred longest path, we constructed a predictive model that estimates the conditional probability *P* (*y*_*t*_ = 1|**x**_<*t*_). Here, *y*_*t*_ ∈ {0, 1} indicates the mutation status of the target gene, and **x**_<*t*_ represents the observed mutation history preceding *t*.

The definition of **x**_<*t*_ determines the scope of the biological information available to the model. We compare two distinct formulations of this input context to test different hypotheses:

#### 1. Path-only context

The input history **x**_<*t*_ is restricted to genes in the longest path that precede *t*. This assumes that the longest path captures the primary mutational trajectory and treats off-path mutations as extraneous information for the prediction task.

#### 2. Full topological context

The input history **x**_<*t*_ is expanded to include all genes that appear before *t* in the global topological ordering of the DAG. This strategy assumes that, although the longest path represents a primary mutational trajectory, off-path genes may still provide conditional information about tumor development before mutation *t*.

### 2.6 Machine learning dataset partitions

For the primary graph-derived and bootstrap-path prediction experiments, models were trained using the curated mutation matrix **M** described above. For each random seed, patient samples were randomly partitioned into an 80% training set and a 20% independent test set. The test set was held out from model fitting and threshold selection and was only used for final performance evaluation. Within the training set, a validation subset was used for decision-threshold tuning, selecting the threshold for each target gene that maximized F1-score on validation predictions. Reported metrics were computed on the independent 20% test set and summarized across 15 random seeds.

### 2.7 Model architectures

We implemented three long-range sequence modeling architectures to capture dependencies within the mutational sequences:

#### 2.7.1 Long Short-Term Memory (LSTM)

The first model was a recurrent neural network (RNN) using LSTM units (Hochreiter and Schmidhuber 1997). The network used a single LSTM layer with *d* = 5 units to capture long-term dependencies in the sparse mutational data. The LSTM dynamics are defined as:

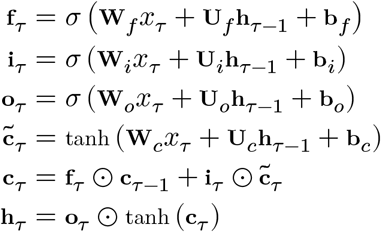

In these equations, τ indexes the position within the input history **x**_<*t*_ for a given target gene *t*, and *x*_τ_ ∈ {0, 1} is the scalar mutation status at position τ. The matrices **W** ∈ ℝ^*d*×1^ are input weights that project the scalar input into the hidden dimension *d*; the matrices **U** ∈ ℝ^*d*×*d*^ are recurrent weights that operate on the previous hidden state **h**_τ−1_; and **b** ∈ ℝ^*d*^ are bias vectors. Each parameter is subscripted by its corresponding gate and learned during training. The symbol *σ* denotes the sigmoid activation function, which outputs values in the range (0, 1), while tanh is the hyperbolic tangent function. The information flow is regulated by three distinct gates: the forget gate **f**_τ_, which determines the extent to which the previous cell state **c**_τ−1_ is retained; the input gate **i**_τ_, which controls how much of the candidate cell state 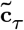 is integrated into the current cell state **c**_τ_ ; and the output gate **o**_τ_, which filters the cell state to create the hidden/output state **h**_τ_. The operator ⊙ represents the Hadamard product. This architecture allows the model to selectively include or forget information about mutation patterns across its input sequence.

#### 2.7.2 Dilated Convolutional Neural Network (CNN)

As an alternative architecture, we implemented a dilated one-dimensional Convolutional Neural Network (CNN). While LSTMs use gated memory to retain information over long spans, dilated CNNs expand their receptive field through dilation, allowing them to incorporate information from distant positions in the sequence without recurrence. Whereas LSTMs address vanishing gradients through gated additive memory updates, dilated CNNs avoid recurrent multiplication through time by using feed-forward paths with short gradient routes. Unlike RNNs, which process sequences step by step, CNNs can process input histories in parallel and can learn hierarchical sequence features through increasing dilation rates.

The architecture consisted of seven one-dimensional convolutional layers with 4 filters, a kernel size of 3, and dilation rates of 1, 2, 4, 8, 16, 32, and 64.

#### 2.7.3 Shared LSTM with Attention and Gene Embeddings

The per-gene LSTM and dilated CNN architectures train an independent model for each target gene *t* in the longest path. While this allows each model to specialize to a target gene, it limits each model to fewer than 1,900 training samples under the 80/20 split. Additionally, genes early in the topological order have very short input sequences **x**_<*t*_, sometimes with only one or two predecessors, providing minimal signal for an independent model. To address these limitations, we propose a shared LSTM architecture in which a single model is trained across all target genes, outlined in Figure 2.

**Figure 2.**
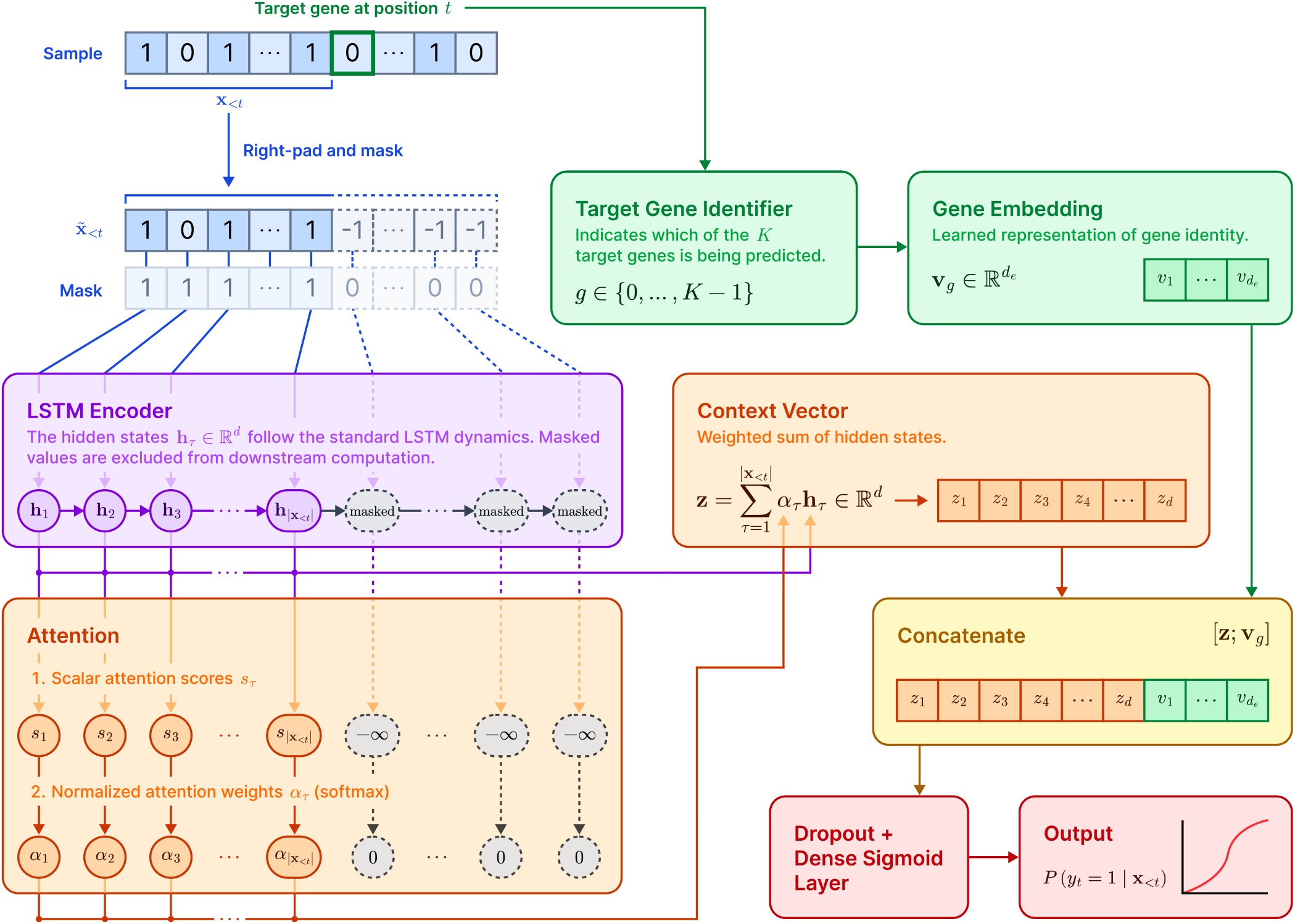
Shared Long Short-Term Memory (LSTM) with attention and gene embeddings for graph-derived mutation prediction. For a given sample, the mutation history preceding target position *t* is extracted as **x**_<*t*_ and right-padded with the sentinel value −1 to form the padded input 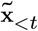. A binary mask distinguishes real predecessor positions from padded positions, so masked entries are ignored by the LSTM encoder and excluded from attention pooling. The LSTM produces hidden states **h**_τ_ for the real input positions, which are scored and normalized into attention weights. The resulting context vector **z** is concatenated with a learned target gene embedding **v**_*g*_ and passed through a dense sigmoid layer to estimate the probability that the target gene is mutated.

The shared model takes two inputs. The first is the predecessor mutation history **x**_<*t*_ ∈ {0, 1}^*T*^, where *T* is the number of real predecessor genes available for target gene *t*. Because |**x**_<*t*_| varies across target genes, this history is right-padded to a common length *T*_max_ before being passed to the model. We denote the padded input as 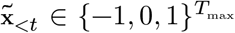, where −1 is a sentinel value used only to mark padded positions. A masking mechanism excludes these sentinel positions from downstream LSTM and attention computations. The second input is an integer gene identifier *g* ∈ {0, …, *K* − 1}, indicating which of the *K* target genes is being predicted.

The padded input 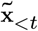 is passed through a masking mechanism and then processed by a single LSTM layer with 16 units, producing a hidden state **h**_τ_ at each position of the original history **x**_<*t*_, where τ = 1, …, |**x**_<*t*_|. The LSTM dynamics follow the same gating equations defined above.

Rather than using only the final hidden state 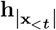, which compresses the predecessor history into a fixed-dimensional representation, we apply an attention pooling mechanism that learns a weighted combination over all positions in **x**_<*t*_ (Yang *et al*. 2016). For each position τ, a scalar attention score is computed:

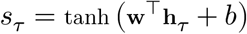

where **w** ∈ ℝ ^*d*^ and *b* ∈ ℝ are learnable parameters. Each hidden state is scored independently, producing a learned measure of how informative that position is for the prediction task. Scores at padded positions are set to −∞ prior to normalization so that they receive zero weight. The scores are normalized via softmax to obtain attention weights:

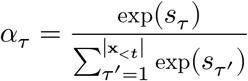

The context vector is then the weighted sum of hidden states:

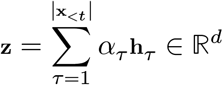

This allows the model to weight informative predecessor positions directly, reducing reliance on the final hidden state as the sole summary of the sequence.

In parallel, the gene identifier *g* is passed through a learned embedding layer that maps each of the *K* target genes into a dense vector 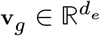, where *d*_*e*_ = 8. These embeddings are initialized randomly and updated during training, allowing the model to learn latent representations of gene identity and gene-specific predictive structure.

The context vector and gene embedding are concatenated and passed through a dropout layer (*p* = 0.3) followed by a single dense unit with sigmoid activation to produce the estimated conditional probability.

Training is performed over all pooled gene–sample pairs from all target genes simultaneously, providing approximately *K* times as many training examples as any single per-gene model. This pooled training, combined with the gene embedding, allows the shared encoder to learn general patterns in upstream mutation structure while the embedding provides gene-specific adaptation.

### 2.8 Training and optimization

Given the sparsity of somatic mutations, the datasets are inherently class imbalanced, with mutations representing rare positive events. To mitigate this, models were trained using binary focal cross-entropy loss (Lin *et al*. 2017). This loss function down-weights easy negatives and focuses learning on hard examples. We optimized the network parameters using the Adam optimizer (Kingma and Ba 2017) for 10 epochs. A batch size of 27 was used on the base LSTM and dilated CNN models, matching Auslander *et al*. (2019), while a batch size of 256 was used on the shared LSTM with attention due to the increased model complexity.

Neural network training is stochastic because of random weight initialization and data shuffling. To assess the consistency of our results, we repeated training and evaluation across 15 independent random seeds using the data partitioning procedure described above. This allowed the reported performance metrics to reflect variations across data splits and initialization states.

For the primary graph-derived experiments, the cooccurrence graph was inferred once from the full mutation matrix **M**, and the resulting weighted and unweighted paths were held fixed across all 15 random seeds. We used this design so that each seed evaluated the same candidate mutational order and prediction tasks, meaning variation across seeds reflects model initialization and train/test partitioning rather than changes in the inferred graph itself. Re-inferring a new graph for each random split would produce different paths across seeds, making the resulting performance estimates less directly comparable.

### 2.9 Evaluation

#### 2.9.1 Metrics

We evaluated mutation-prediction performance using standard binary classification metrics calculated from the confusion matrix: recall, the ability of the model to identify true mutations; precision, which measures the proportion of predicted mutations that were actually mutated; F1-score, the harmonic mean of precision and recall; and area under the ROC curve (AUC), a threshold-independent measure of the model’s ability to rank positive samples above negative ones.

#### 2.9.2 Thresholding

To convert predicted probabilities into binary class labels, we employed two strategies. The first strategy used a fixed threshold of 0.5: predicted probabilities in [0.5, 1] were labeled as mutations, and probabilities in [0, 0.5) were labeled as non-mutations. The second approach tuned this threshold separately for each model. For each gene *t*, we performed a grid search from 0.25 to 0.95 on the validation subset of the training data to select the threshold that maximized the F1-score for that target gene. The selected threshold was then applied to the independent test set. Optimizing F1-score balances missed mutations and false positives. In contrast, recall can be increased trivially by lowering the threshold, while precision can often be increased by raising the threshold and making fewer positive predictions. We report results for both fixed and tuned/adaptive strategies across all considered paths and models.

## 3 Results

### 3.1 Clusters and stages

No clustering method recovered AJCC stage from the mutation data. On the full gene set, Jaccard hierarchical clustering, fixed at four clusters, produced a highly imbalanced result where the largest cluster contained 82% of samples, with Stage II dominant in all but one cluster (ARI = −0.009). Louvain and Leiden community detection identified 12 and 15 communities respectively, each with a stage distribution similar to that of the full sample set (Louvain ARI = 0.005, Leiden ARI = 0.005). Ward hierarchical clustering in PCA space selected eight clusters via silhouette score, though ARI was still rather poor at -0.013. Filtering to the set of genes mutated in ≥ 5% of samples did not improve these results. The first principal component was near-perfectly correlated with tumor mutational burden (Pearson *r* = 0.999 unfiltered, *r* = 0.974 filtered), which confirms that the dominant axis of variation reflects mutational load instead of stage-specific or molecular patterns.

### 3.2 Baselines

To establish a comparative baseline, we replicated the mutation sequence modeling experiment from Auslander *et al*. (2019). We trained LSTM models using the training cohort defined in their study and evaluated performance on the dataset from Giannakis *et al*. (2016) using the same fixed global gene ordering. These replicated results (Tab. 1) provide a baseline for evaluating the proposed graph-derived ordering strategies. Because the dataset is heavily imbalanced toward non-mutated labels, a random Bernoulli classifier matched to the empirical class distribution would achieve an expected accuracy of approximately 0.96 but a precision equal to the mutation prevalence, approximately 0.02. This highlights the inadequacy of accuracy, and accuracy-like metrics, under extreme class imbalance.

### 3.3 Inference of evolutionary trajectories

To map the dependency structure of colorectal adenocarcinoma, we reconstructed the mutational landscape as a directed acyclic graph and identified dominant evolutionary trajectories using both unweighted and weighted path-finding algorithms. The resulting graph, shown in part in Figure 3, shows that maximizing sequence length (unweighted) and maximizing cumulative edge weight (weighted) yield distinct evolutionary hypotheses. This divergence is illustrated in Figure 4, which contrasts the prevalence-heavy profile of the unweighted path with the profile of the weighted path. We then evaluated these inferred trajectories through downstream mutation prediction.

**Figure 3.**
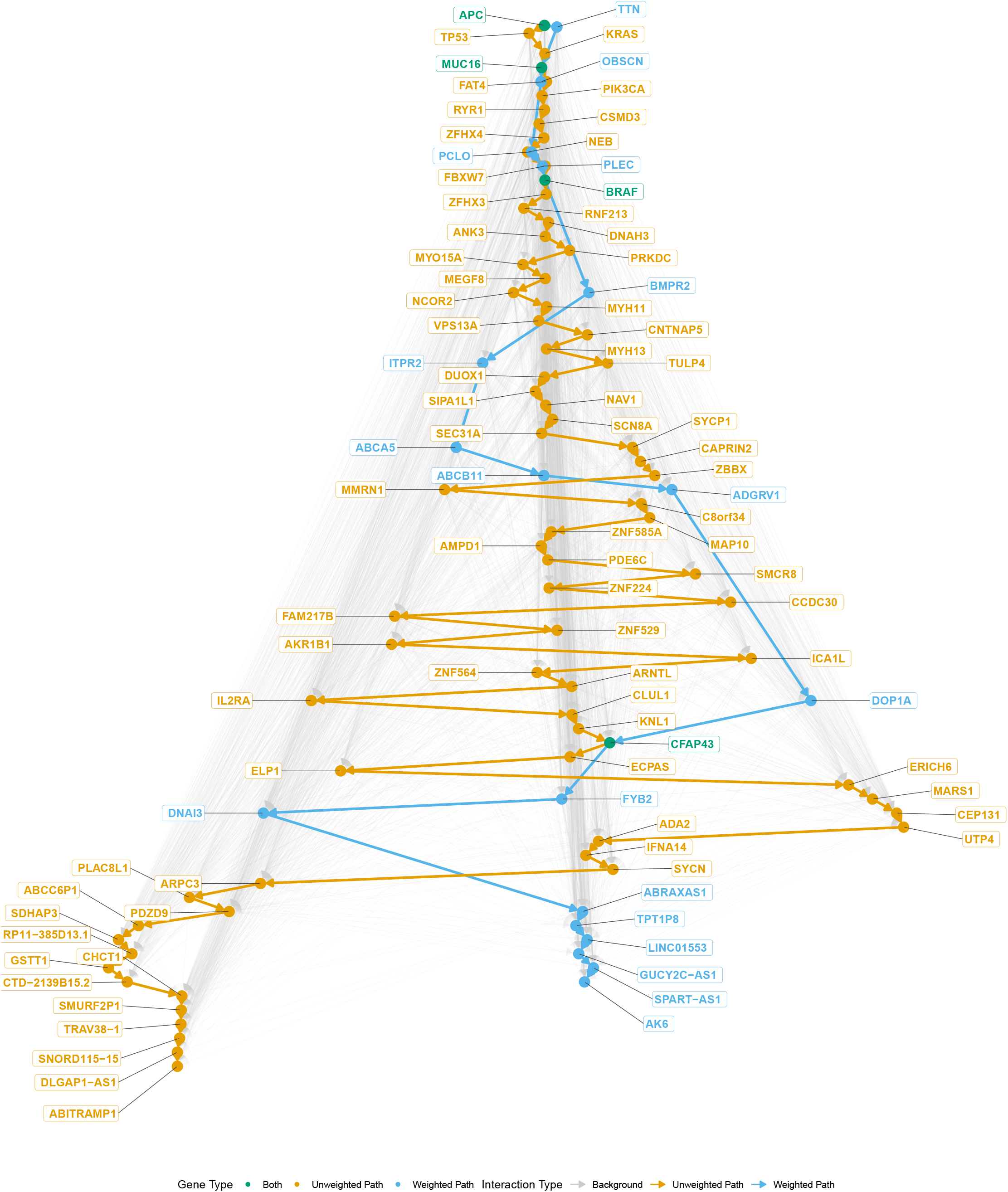
Weighted and unweighted longest mutational paths. The directed acyclic graph (DAG) is arranged using a Sugiyama layout to reflect topological order (Sugiyama *et al*. 1981), with earlier inferred mutational events at the top and later inferred events at the bottom. Nodes and edges are colored by their inclusion in the computed longest paths. Green indicates genes identified by both the weighted and unweighted paths; orange denotes the unweighted longest path, maximizing vertex count; and blue denotes the weighted longest path, maximizing the cumulative edge weight.

**Figure 4.**
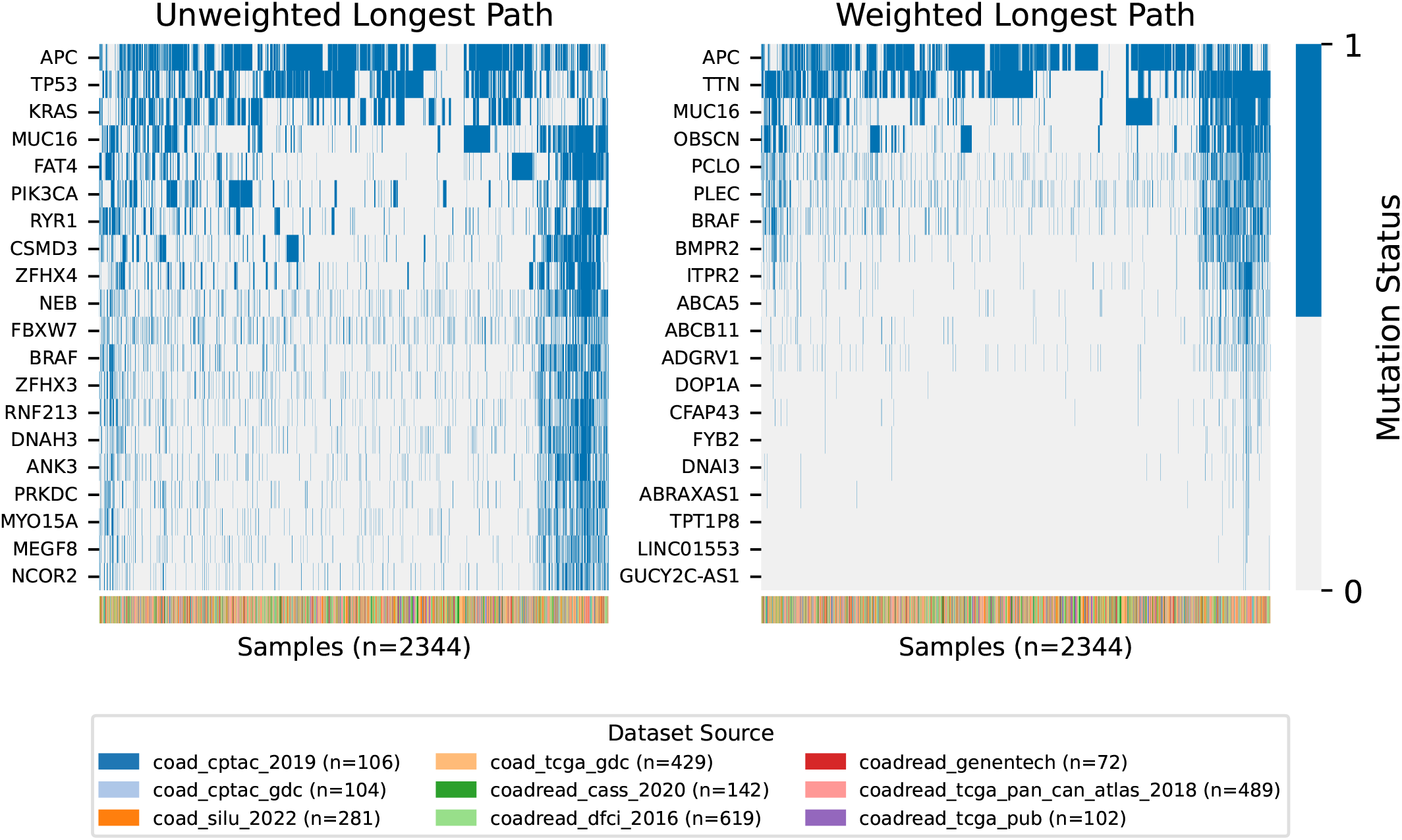
Binary mutation statuses (*m*_*ik*_) for the genes comprising the inferred unweighted (left) and weighted (right) longest paths across the 2,344 samples. Rows correspond to the first 20 genes included in each trajectory, while columns represent patient samples ordered using hierarchical clustering (Euclidean distance, average linkage). Blue indicates the presence of a mutation, and light gray indicates the absence of a mutation. The color-coded annotation bar at the bottom of each heatmap identifies the originating dataset for each sample.

### 3.2 Predictive performance of sequence models

We used the inferred paths to train LSTM models, dilated CNN models, and shared LSTM models with attention and gene embeddings to predict the presence of a mutation at gene *t*, given the patient’s prior mutational history **x**_<*t*_. Table 2 summarizes predictive performance across path construction strategies for the standard LSTM architecture.

**Table 1.** Baseline confusion matrix and metrics.

| Actual | Predicted |  | Total |
| --- | --- | --- | --- |
|  | Positive | Negative |  |
| Positive | 438 | 147,956 | 148,394 |
| Negative | 1,133 | 7,478,410 | 7,479,543 |
| Total | 1,571 | 7,626,366 | 7,627,937 |
| Recall |  |  | 0.003 |
| Precision |  |  | 0.279 |
| Accuracy |  |  | 0.980 |
| F1-score |  |  | 0.006 |

**Table 2.** Performance metrics (mean ± SD over *n* = 15 random seeds) for LSTM models using fixed (0.5) and tuned thresholds across contexts and path strategies.

| Configuration | Recall | Precision | AUC | F1-score |
| --- | --- | --- | --- | --- |
| **Unweighted Longest Path** |  |  |  |  |
| Path-only (Fixed) | 0.201 ± 0.018 | **0.608 ± 0.027** | 0.902 ± 0.003 | 0.302 ± 0.019 |
| Path-only (Tuned) | 0.643 ± 0.029 | 0.383 ± 0.015 | 0.902 ± 0.003 | 0.479 ± 0.014 |
| Full context (Fixed) | 0.265 ± 0.020 | 0.604 ± 0.020 | 0.897 ± 0.002 | 0.368 ± 0.019 |
| Full context (Tuned) | 0.646 ± 0.016 | 0.399 ± 0.029 | 0.897 ± 0.002 | 0.493 ± 0.023 |
| **Weighted Longest Path** |  |  |  |  |
| Path-only (Fixed) | 0.314 ± 0.025 | 0.552 ± 0.023 | **0.914 ± 0.004** | 0.400 ± 0.025 |
| Path-only (Tuned) | 0.699 ± 0.027 | 0.439 ± 0.022 | **0.914 ± 0.004** | 0.539 ± 0.014 |
| Full context (Fixed) | 0.441 ± 0.021 | 0.576 ± 0.018 | **0.914 ± 0.005** | 0.499 ± 0.016 |
| Full context (Tuned) | **0.749 ± 0.028** | 0.443 ± 0.020 | **0.914 ± 0.005** | **0.556 ± 0.016** |

Note: AUC is threshold-independent; each AUC value applies to both the fixed- and tuned-threshold rows for the same context. Bold values indicate the best mean performance within each metric column.

Despite excluding *TP53*, the weighted longest path outperformed the unweighted approach across most metrics. The weighted model achieved a maximum AUC of 0.914, compared to 0.902 for the unweighted model. This suggests that the weighted sequence, despite omitting a canonical driver, provides a stronger predictive signal than the unweighted chain. Several graph-derived configurations improved precision over the reproduced baseline of 0.279, particularly under fixed-threshold evaluation.

When comparing architectures, the LSTM and dilated CNN achieved broadly comparable performance; dilated CNN results are reported in Table 3. Dilated CNN achieved competitive recall on the tuned full-context path (0.747 vs 0.749 for LSTM), though it showed lower recall (0.329 vs 0.441) in the fixed threshold, full-context configuration. In addition, both architectures identified the fixed-threshold, full-context models as having the highest precision among the tested configurations, suggesting broad agreement across evaluation metrics.

**Table 3.** Performance metrics (mean ± SD over *n* = 15 random seeds) for dilated CNN models using fixed (0.5) and tuned thresholds across contexts.

| Configuration | Recall | Precision | AUC | F1-score |
| --- | --- | --- | --- | --- |
| **Unweighted Longest Path** |  |  |  |  |
| Path-only (Fixed) | 0.148 ± 0.050 | **0.541 ± 0.059** | 0.851 ± 0.034 | 0.227 ± 0.062 |
| Path-only (Tuned) | 0.608 ± 0.042 | 0.292 ± 0.019 | 0.851 ± 0.034 | 0.393 ± 0.017 |
| Full context (Fixed) | 0.225 ± 0.049 | 0.484 ± 0.101 | 0.860 ± 0.035 | 0.304 ± 0.058 |
| Full context (Tuned) | 0.632 ± 0.040 | 0.282 ± 0.037 | 0.860 ± 0.035 | 0.388 ± 0.033 |
| **Weighted Longest Path** |  |  |  |  |
| Path-only (Fixed) | 0.337 ± 0.081 | 0.485 ± 0.067 | 0.859 ± 0.074 | 0.391 ± 0.061 |
| Path-only (Tuned) | 0.691 ± 0.085 | 0.362 ± 0.024 | 0.859 ± 0.074 | **0.473 ± 0.031** |
| Full context (Fixed) | 0.329 ± 0.107 | 0.539 ± 0.085 | **0.893 ± 0.019** | 0.391 ± 0.091 |
| Full context (Tuned) | **0.747 ± 0.031** | 0.327 ± 0.051 | **0.893 ± 0.019** | 0.452 ± 0.052 |

Note: AUC is threshold-independent; each AUC value applies to both the fixed- and tuned-threshold rows for the same context. Bold values indicate the best mean performance within each metric column.

The proposed shared LSTM with attention and gene embeddings followed the same overall trends as the simpler models. Compared with the standard LSTM, the shared LSTM showed lower recall but similar precision under fixed-threshold evaluation. Under tuned-threshold evaluation, recall was higher than that of the standard LSTM (0.804 on the weighted, full-context, tuned configuration vs. 0.749 on the same configuration for standard LSTM), though precision is often comparable or lower. Although the shared attention model did not uniformly outperform the standard LSTM across all metrics, it achieved competitive AUC and high recall while providing a single gene-conditioned model across all targets.

### 3.5 Impact of context and thresholding

Incorporating off-path information from the background topological order improved predictive performance. Models trained with full topological context, which includes all predecessor genes in the DAG rather than only genes in the longest path, achieved the highest performance. Specifically, under the weighted strategy, adding full context improved the F1-score from 0.539 to 0.556 and the recall from 0.699 to 0.749 (Table 2). This suggests that off-path mutations contain information that can refine prediction of downstream mutational states.

Threshold tuning was also important for handling the inherent class imbalance of somatic mutation data. Standard fixed thresholds (0.5) frequently failed to capture the minority positive class. Tuning the decision threshold to maximize F1-score on a validation set substantially improved recall, reaching 0.749 for the weighted, full-context LSTM configuration.

However, this gain in recall came with a trade-off in precision compared with fixed-threshold evaluation (e.g., dropping from 0.576 to 0.443 in the weighted, full context configuration). This reduction occurs because optimizing for F1-score in sparse datasets requires lowering the decision boundary to capture rare mutational events. Although this increases the false positive rate, the substantial recovery in recall may justify the reduction in precision for mutation screening tasks.

### 3.6 Bootstrapping results

For the bootstrap consensus graph, we obtained both weighted and unweighted longest paths. These paths were then used to train models with the standard LSTM architecture; evaluation metrics for those models are reported Note: AUC is threshold-independent; each AUC value applies to both the fixed- and tuned-threshold rows for the same context. Bold values indicate the best mean performance within each metric column. in Table 5. Unlike primary graph results, the unweighted bootstrap path performed slightly better than the weighted path on most metrics. In addition, full topological context did not improve performance relative to the path-only context.

**Table 4.**
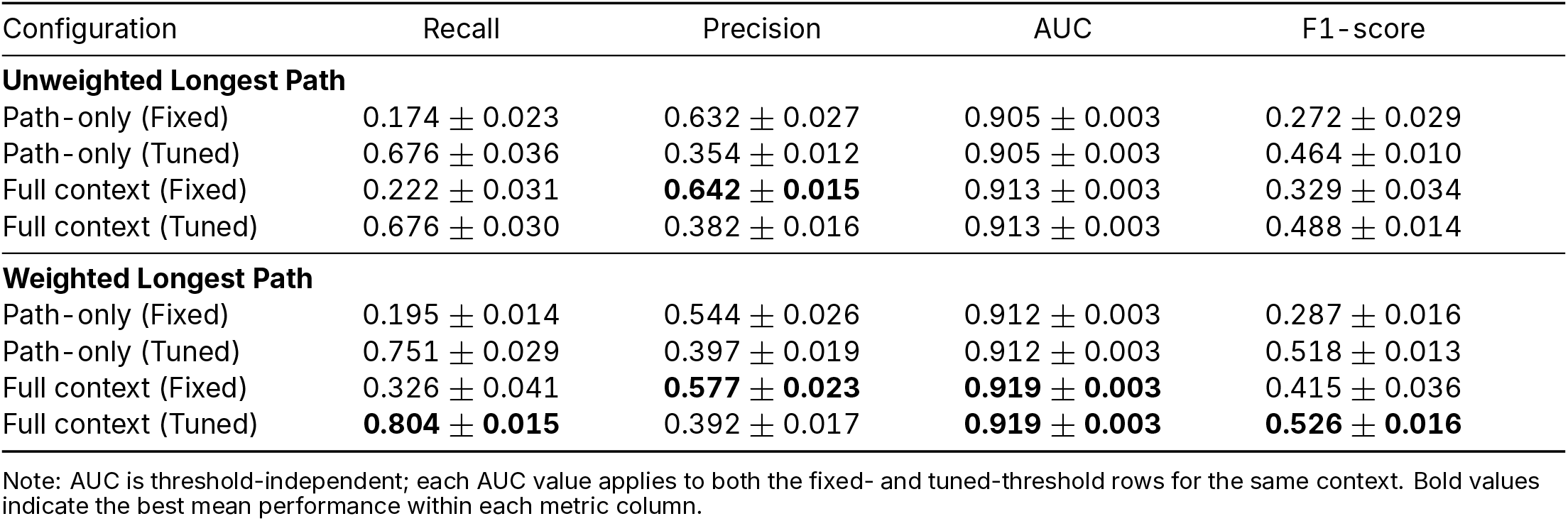
Performance metrics (mean ± SD over *n* = 15 random seeds) for shared LSTM models with attention using fixed (0.5) and tuned thresholds across contexts and path strategies.

**Table 5.** Performance metrics (mean ± SD over *n* = 15 random seeds) for the standard LSTM architecture on the bootstrap consensus paths using fixed (0.5) and tuned thresholds across contexts and path strategies.

| Configuration | Recall | Precision | AUC | F1-score |
| --- | --- | --- | --- | --- |
| **Unweighted Longest Path** |  |  |  |  |
| Path-only (Fixed) | 0.192 ± 0.016 | 0.592 ± 0.018 | **0.840 ± 0.005** | 0.289 ± 0.020 |
| Path-only (Tuned) | **0.580 ± 0.025** | 0.362 ± 0.017 | **0.840 ± 0.005** | 0.445 ± 0.017 |
| Full context (Fixed) | 0.201 ± 0.013 | 0.586 ± 0.021 | 0.836 ± 0.005 | 0.299 ± 0.017 |
| Full context (Tuned) | **0.580 ± 0.024** | 0.365 ± 0.017 | 0.836 ± 0.005 | **0.448 ± 0.017** |
| **Weighted Longest Path** |  |  |  |  |
| Path-only (Fixed) | 0.154 ± 0.016 | **0.610 ± 0.025** | 0.836 ± 0.005 | 0.246 ± 0.021 |
| Path-only (Tuned) | 0.563 ± 0.028 | 0.363 ± 0.016 | 0.836 ± 0.005 | 0.441 ± 0.017 |
| Full context (Fixed) | 0.164 ± 0.016 | 0.606 ± 0.027 | 0.831 ± 0.005 | 0.258 ± 0.022 |
| Full context (Tuned) | 0.561 ± 0.026 | 0.365 ± 0.017 | 0.831 ± 0.005 | 0.442 ± 0.017 |

Note: AUC is threshold-independent; each AUC value applies to both the fixed- and tuned-threshold rows for the same context. Bold values indicate the best mean performance within each metric column.

The lower performance of the bootstrap consensus paths compared with the primary graph-derived paths suggests that using a smaller data cohort for graph inference reduces the predictive quality of the resulting order, but the above-baseline performance indicates that the signal is not entirely dependent on constructing the graph from the full matrix.

## 4 Discussion

The unweighted longest path derived from the primary graph construction method recovered several established colorectal cancer drivers, including *APC, TP53, KRAS*, and *BRAF*, among its early vertices and contained 75 genes in total. However, the inferred ordering deviated from standard linear progression models, with *TP53* appearing before *KRAS* in the sequence *APC* → *TP53* → *KRAS* (Vogelstein *et al*. 2013). This suggests that while the unweighted approach captures high-prevalence drivers, maximizing path length may favor placing high-degree vertices earlier in the path to extend the chain.

In contrast, the weighted longest path optimized the strength of conditional dependencies rather than the number of vertices, and included 22 genes. Notably, this probabilistic trajectory excluded *TP53*. Rather than forcing a route through all canonical drivers, the weighted algorithm favored a statistically cohesive chain of interactions with stronger pairwise asymmetry. This distinction is consistent with the idea that colorectal tumorigenesis follows a preferred but not strictly fixed order, and that the accumulation of key alterations may be as important as their exact sequence (Fearon and Vogelstein 1990).

Notably, both paths start with *APC*, a canonical initiating driver of the adenoma–carcinoma sequence. Both paths also identify *BRAF* early, consistent with its known role as a colorectal cancer driver. However, the inferred paths may also reflect gene length and mutational opportunity, since long genes such as *TTN* are more likely to accumulate mutations. This is an important limitation of co-occurrence-based inference and suggests that future graph construction should account for gene length, background mutation rate, or sample-level mutational burden.

The paths obtained from the bootstrap consensus graph showed broadly similar trends. The unweighted bootstrap path contained 158 genes and included *APC, TP53, KRAS*, and *BRAF*, while the weighted bootstrap path contained 153 genes and again excluded *TP53*. These results suggest that several core ordering signals are recoverable even when the graph is inferred from a smaller cohort.

The shared LSTM with attention and gene embeddings represents the main predictive architecture proposed in this study. Compared with the standard LSTM and dilated CNN, which train separate models for each target gene, the shared attention model pools information across all target genes while conditioning each prediction on a learned gene embedding. This design is well suited to sparse mutation matrices because many target genes have few positive examples, making independent per-gene models data-limited. Although this architecture did not dominate every metric, its competitive AUC and high recall demonstrate its promise.

Several limitations should be noted. First, a key assumption of this pseudotemporal approach is that inter-patient heterogeneity contains information about different points along mutational progression. The availability of AJCC Stage I–IV samples provides some amount of coverage across disease extent. However, our clinical-stage analysis showed that mutation profiles did not cluster by AJCC stage, while the dominant axis of variation was strongly associated with mutational burden. As such, the inferred ordering likely reflects mutation accumulation and co-occurrence structure more than clinical stage progression. Consistent with this interpretation, supplementary stratified analyses showed that stage- and Louvain-specific graph-derived models gave mixed results.

The fixed-graph design also affects how the predictive results should be interpreted. Holding the primary graph-derived paths constant across random seeds made the seed-level comparisons consistent, since each run evaluated the same target genes and predecessor histories. However, this design does not test how stable the inferred graph would be if it were reconstructed within each training split. The bootstrap consensus analysis provides a partial check by evaluating paths from the smaller COAD cohort used by Auslander *et al*. (2019); these paths performed worse than the primary graph-derived paths but remained above the reproduced baseline. This difference may reflect the smaller cohort used for graph inference, the bootstrap aggregation procedure, the frequency-thresholding strategy used to obtain a DAG, or a combination of these factors. Future work should directly evaluate graph stability with training-split specific inference.

In conclusion, this study demonstrates the potential of combining graph-based mutational ordering with long-term sequence models for mutation prediction in colon cancer. Future work should evaluate additional branching paths from the co-occurrence asymmetry graph and further validate inferred gene orders using long-term learning models.

## Author contributions

Collin Sumrell (Conceptualization [supporting], Software [lead], Methodology [supporting], Formal analysis [equal], Investigation [equal], Data curation [equal], Validation [equal], Visualization [lead], Writing-original draft [equal], Writing-review & editing [equal]), Aleksandr Shishkin (Data curation [supporting], Writing-review & editing [supporting]), Ekene Okeke (Software [supporting], Formal analysis [supporting]), Alexander Zelikovsky (Formal analysis [supporting], Investigation [supporting], Supervision [supporting]), Marmar R. Moussa (Conceptualization [lead], Methodology [lead], Formal analysis [equal], Investigation [equal], Project administration [lead], Software [supporting], Validation [equal], Writing-original draft [equal], Writing-review & editing [equal], Supervision [lead], Funding acquisition [lead])

## Supplementary material

Supplementary material is available as a separate file accompanying this manuscript.

## Data availability

The data and code underlying this study are available at github.com/moussa-lab/MutationPrediction.

