## Supplementary material for "Predicting Gene Mutations in Colon Cancer Using Long-Term Temporal Dependency Learning on a Directed Co-Occurrence Asymmetry Graph": SumrellMutationPrediction_supplement.pdf


**Key Points**

- Mutation-profile clusters did not recover AJCC clinical stage
- Stage- and cluster-stratified prediction experiments showed mixed performance and should be interpreted as exploratory rather than primary evidence

### S1 Supplementary methods

#### S1.1 Data used for clustering and stratified prediction

We performed supplementary analyses to evaluate whether the proposed graph-derived mutation prediction framework behaved differently across clinical stage subsets or mutation-profile-derived clusters. These analyses used the same curated mutation matrix described in the main text (Li *et al.* 2020; Cancer Genome Atlas Network 2012; Heath *et al.* 2021; Roelands *et al.* 2023; Vasaikar *et al.* 2019; Giannakis *et al.* 2016; Seshagiri *et al.* 2012; Cerami *et al.* 2012; Gao *et al.* 2013; de Bruijn *et al.* 2023). For analyses involving clinical stage, we restricted the cohort to the 1,516 samples with available AJCC stage annotations. These included 246 Stage I, 552 Stage II, 452 Stage III, and 266 Stage IV tumors. Samples with unknown or missing stage annotations were excluded from stage comparison analyses.

Tumor mutational burden was computed for each sample. To evaluate whether rare mutation events influenced clustering structure, clustering analyses were performed on both the full mutation matrix and a filtered matrix retaining genes mutated in at least 5% of samples. The filtered matrix contained 1,453 genes. Metrics may vary slightly from results reported in the paper due to differences in random seeds.

#### S1.2 Unsupervised clustering of staged samples

We evaluated whether unsupervised clustering of binary somatic mutation profiles recovered AJCC clinical stage (Amin *et al.* 2017). Principal component analysis (PCA) was performed on the staged samples, and the elbow of the scree plot selected three principal components for downstream graph-based community detection. We also computed the Pearson correlation between the first principal component and tumor mutational burden .

Four clustering strategies were evaluated. First, we performed hierarchical clustering using Jaccard distance on binary mutation profiles and Ward linkage (Ward 1963), with the number of clusters fixed at $k = 4$ to match the four AJCC stages. Second, we constructed a $k$-nearest neighbor graph in three-dimensional PCA space using $k = 15$ neighbors and applied Louvain community detection (Blondel *et al.* 2008). Third, we applied Leiden community detection to the same PCA-derived $k$NN graph (Traag *et al.* 2019). Foruth, we performed Ward hierarchical clustering directly in PCA space, selecting $k$ by silhouette score.

Cluster-stage agreement was evaluated using adjusted Rand index (ARI) and normalized mutual information (NMI) (Hubert and Arabie 1985; Strehl and Ghosh 2003). ARI measures agreement between two partitions after correcting for chance, with values near zero indicating random agreement. NMI measures shared information between the inferred clusters and stage labels.

#### S1.3 Stage-stratified graph construction and prediction

We performed exploratory stage-stratified prediction experiments to test whether graph-derived mutation prediction differed across AJCC stages or was notably different from stage-agnostic, full-cohort inference and prediction. For each stage, a separate directed co-occurrence asymmetry graph was constructed using only samples from that stage. A weighted longest path was then extracted from each stage-specific graph. We focused on the weighted path for the stage-stratified experiments because it was the best-performing path strategy in the primary full-cohort analysis.

Standard LSTM models were trained and evaluated separately within each stage using the same prediction task formulation described in the main text. Both path-only and full-context input definitions were evaluated. For each stage and context, model performance was summarized over 15 random seeds. We report results using both a fixed decision threshold of 0.5 and an adaptive threshold selected to maximize F1-score on validation predictions.

#### S1.4 Louvain-cluster-stratified graph construction and prediction

Because mutation-profile clusters did not correspond to AJCC stage, we also evaluated prediction within the two largest Louvain communities. For each of these two communities, a separate co-occurrence asymmetry graph was constructed. Both weighted and unweighted paths were extracted from each cluster-specific graph.

Standard LSTM models were trained and evaluated within each Louvain community using both path-only and full-context input definitions. Results were summarized over 5 random seeds. As in the stage-stratified analysis, we report both fixed-threshold and F1-tuned-threshold performance. Because these subgroup analyses used smaller datasets than the primary full-cohort analysis, the stratified prediction results were treated as exploratory analyses rather than primary evidence.

### S2 Supplementary results

#### S2.1 Mutation profile clusters do not recover AJCC stage

Across all clustering approaches, unsupervised mutation-profile clusters showed little agreement with AJCC clinical stage. In the filtered 1,453-gene matrix, Jaccard hierarchical clustering with Ward linkage produced a highly imbalanced four-cluster solution, with one cluster containing most staged samples (Supplementary Figure S1). Agreement with stage was near random (ARI $= -0.013$, NMI $= 0.028$). The row normalized heatmap in Supplementary Figure S1 shows that Stage II was the dominant stage in all four clusters.

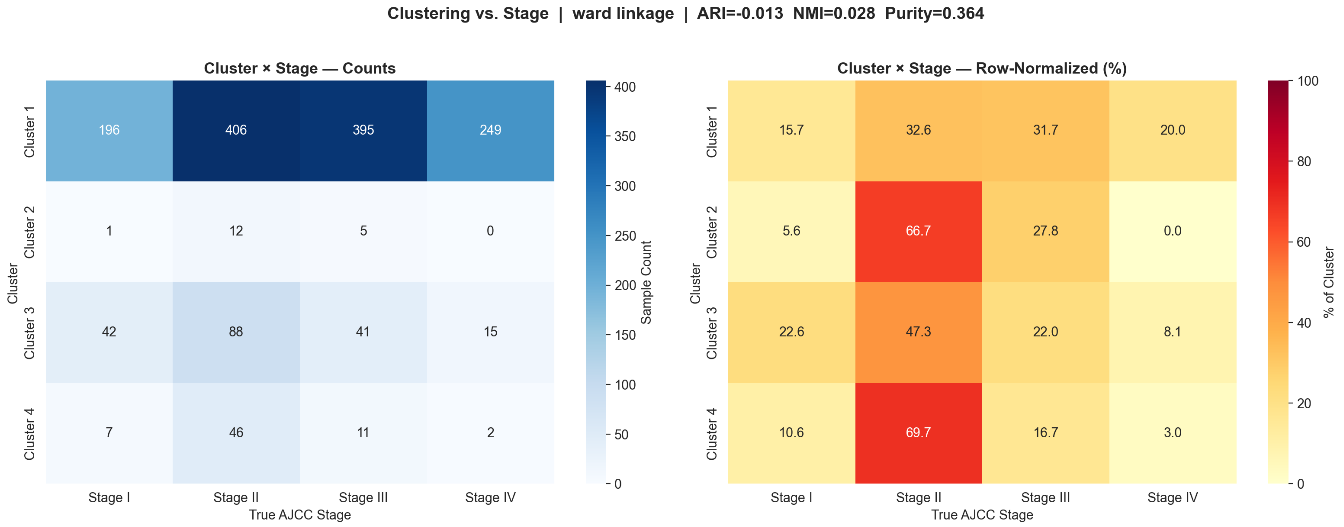


**Supplementary Figure S1** Jaccard hierarchical clustering versus AJCC stage using Ward linkage and $k = 4$ on the filtered mutation matrix. The left heatmap shows cluster-by-stage counts and the right heatmap shows row-normalized samples.

The PCA visualizations support the same conclusion. In Supplementary Figure S2, samples separate primarily along PC1 when colored by Ward cluster assignment, but the same projection colored by AJCC stage shows extensive mixing of

Stage I–IV samples. PC1 was strongly correlated with tumor mutational burden ($r = 0.974$) in the filtered matrix, indicating that the dominant axis of variation reflected mutational burden rather than clinical stage.

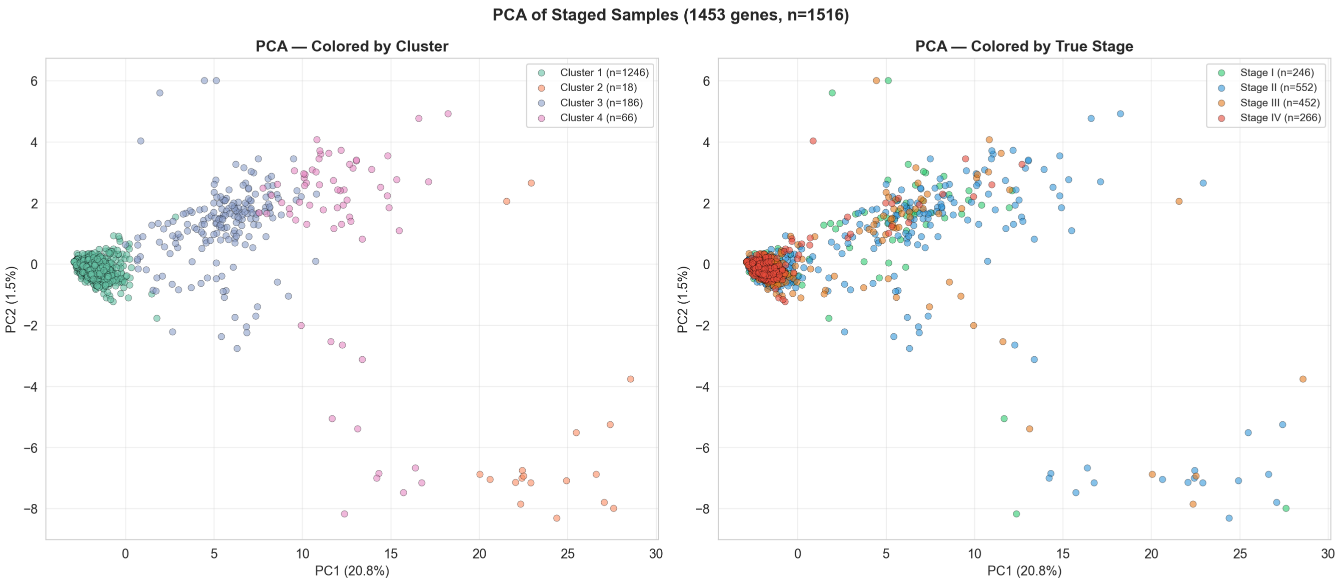


**Supplementary Figure S2** PCA of staged samples after filtering to genes mutated in at least 5% of samples. Points are clustered by Ward cluster assignment (left) and true AJCC stage (right).

Community detection in PCA space produced more balanced partitions than Jaccard hierarchical clustering, but still did not recover clinical stage. Louvain community detection identified 12 communities with ARI $= 0.005$ and NMI $= 0.019$ (Supplementary Figures S3 and S4). The PCA projection in Supplementary Figure S3 shows that Louvain communities partition mutation-profile space but fails to replicate stage intermixing across this space. The cluster-by-stage heatmaps in Supplementary Figure S4 show that every Louvain cluster contrains a mix of AJCC stages.

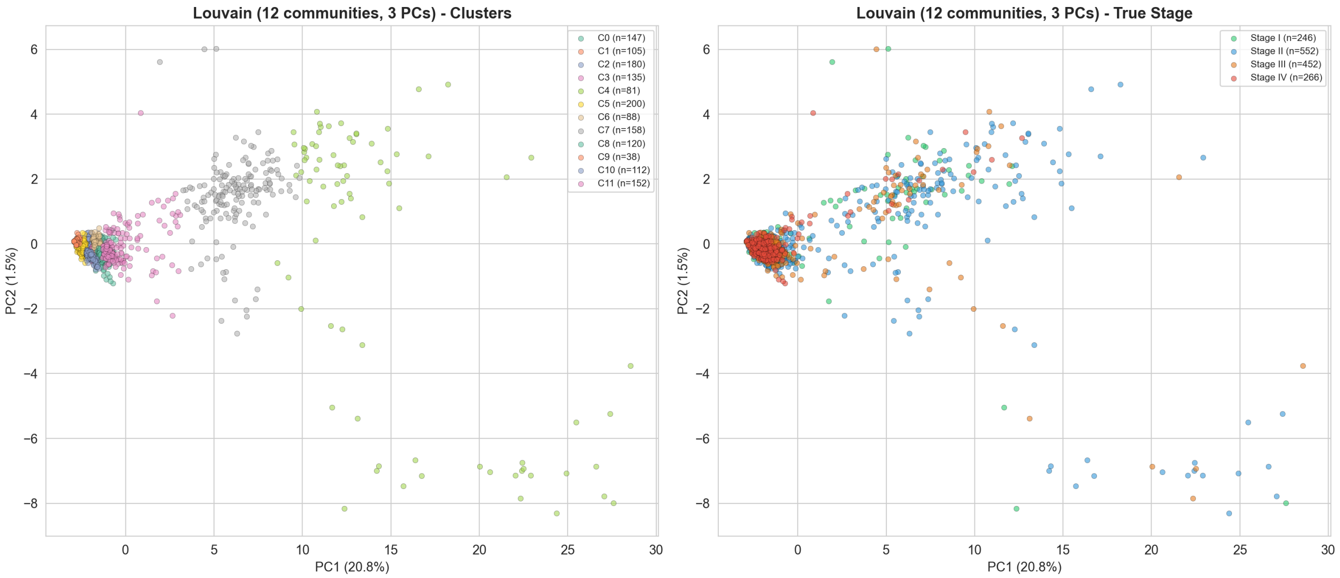


**Supplementary Figure S3** PCA of staged samples after filtering to genes mutated in at least 5% of samples. Points are clustered by Louvain community assignment (left) and true AJCC stage (right).

Leiden community detection produced the same qualitative result, with 13 communities and ARI $= 0.006$, NMI $= 0.020$ (Supplementary Figure S5). Ward clustering in PCA space selected $k = 3$ by silhouette score but again failed to recover clinical stage, with ARI $= -0.009$ and NMI $= 0.026$ (Supplementary Figure S6)

Repeating the full analysis without the 5% mutation-frequency filter changed little. Together, these results indicate that unsupervised clustering of binary somatic mutation profiles did not recover AJCC clinical stage in this cohort.

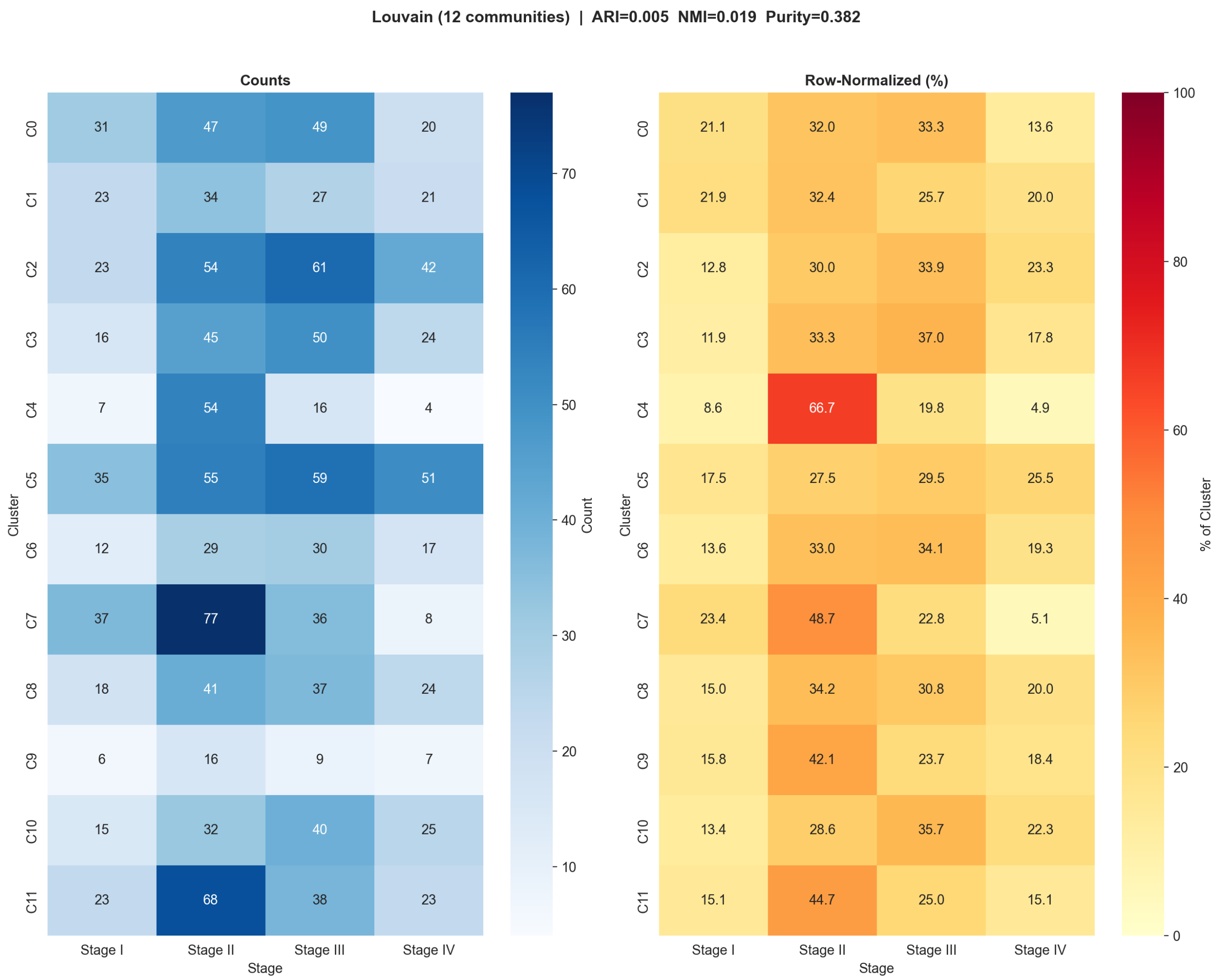


**Supplementary Figure S4** Louvain community detection versus AJCC stage on the filtered mutation matrix. The left heatmap shows cluster-by-stage counts and the right heatmap shows row-normalized samples.

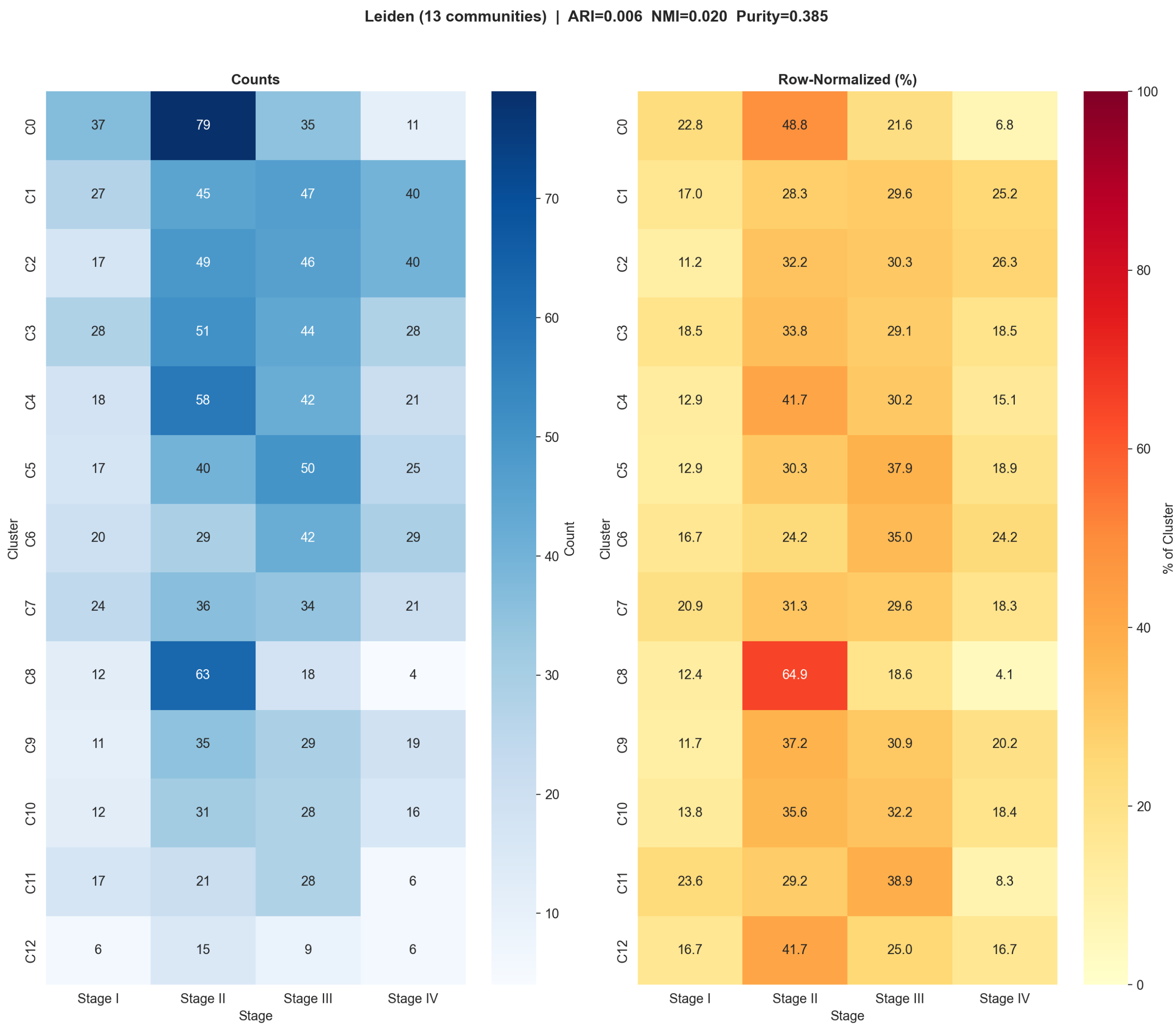


**Supplementary Figure S5** Leiden community detection versus AJCC stage on the filtered mutation matrix. The left heatmap shows cluster-by-stage counts and the right heatmap shows row-normalized samples.

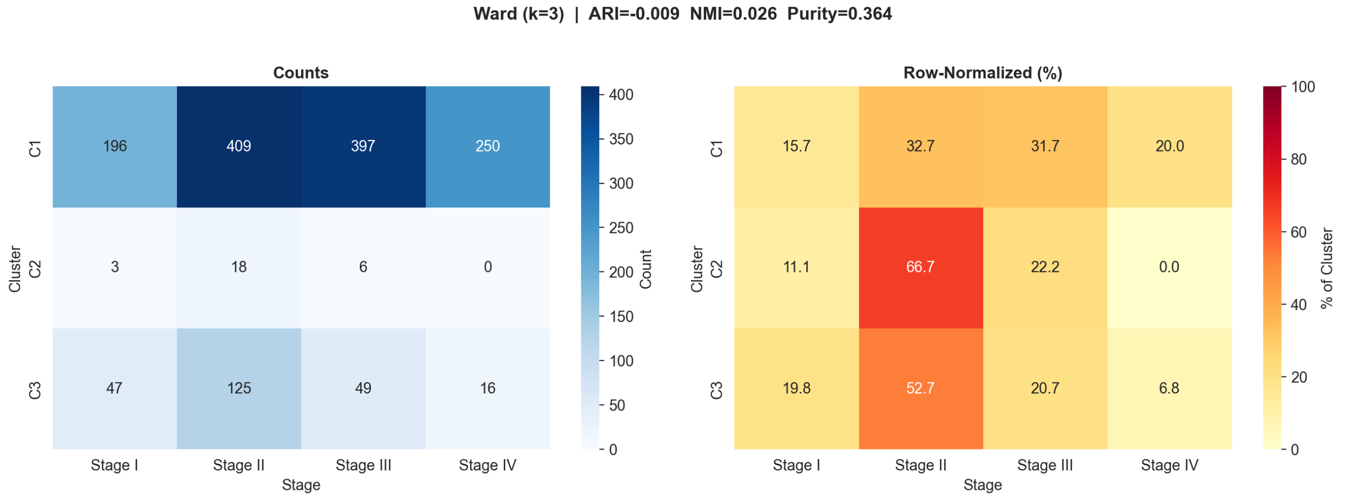


**Supplementary Figure S6** Ward clustering using $k = 3$ samples, with $k$ selected by silhouette score, versus AJCC stage on the filtered mutation matrix. The left heatmap shows cluster-by-stage counts and the right heatmap shows row-normalized samples.

#### S2.2 Stage-stratified prediction results

Stage-stratified graph-derived prediction results are reported in Supplementary Table S1. Performance varied across AJCC stages, with Stage II and Stage III generally showing stronger tuned-threshold AUC and F1-score than Stage I and Stage IV. However, F1-based threshold tuning did not consistently improve test-set F1-score. In Stage I, Stage III, and Stage IV, tuned-threshold models often showed much higher recall but lower precision and lower test-set F1-score than the corresponding fixed-threshold models.

One possible explanation is that the smaller stage-specific cohorts make validation-based threshold selection less stable. Because the adaptive threshold is selected on a validation subset within the training data, small or unusually distributed validation splits may favor thresholds that maximize validation F1 but do not transfer well to the independent test split. We therefore treat the tuned-threshold stage-stratified results cautiously and report these experiments as exploratory supplementary analyses.

**Supplementary Table S1** Stage-stratified standard LSTM performance using weighted longest paths. Values are mean $\pm$ SD over 15 random seeds.

| Stage | Context | Threshold | Recall | Precision | AUC | F1 |
|---|---|---|---|---|---|---|
| I | Path-only | Fixed | $0.370 \pm 0.081$ | $0.510 \pm 0.085$ | $0.690 \pm 0.020$ | $0.420 \pm 0.060$ |
| I | Path-only | Tuned | $\mathbf{0.959 \pm 0.033}$ | $0.122 \pm 0.027$ | $0.690 \pm 0.020$ | $0.215 \pm 0.042$ |
| I | Full | Fixed | $0.335 \pm 0.078$ | $0.546 \pm 0.071$ | $0.727 \pm 0.031$ | $0.408 \pm 0.067$ |
| I | Full | Tuned | $0.880 \pm 0.076$ | $0.116 \pm 0.025$ | $0.727 \pm 0.031$ | $0.204 \pm 0.038$ |
| II | Path-only | Fixed | $0.283 \pm 0.054$ | $0.554 \pm 0.068$ | $0.892 \pm 0.014$ | $0.370 \pm 0.052$ |
| II | Path-only | Tuned | $0.884 \pm 0.050$ | $0.268 \pm 0.032$ | $0.892 \pm 0.014$ | $0.410 \pm 0.039$ |
| II | Full | Fixed | $0.383 \pm 0.046$ | $0.611 \pm 0.060$ | $\mathbf{0.917 \pm 0.015}$ | $0.467 \pm 0.034$ |
| II | Full | Tuned | $0.780 \pm 0.033$ | $0.458 \pm 0.093$ | $\mathbf{0.917 \pm 0.015}$ | $\mathbf{0.571 \pm 0.076}$ |
| III | Path-only | Fixed | $0.540 \pm 0.084$ | $0.564 \pm 0.043$ | $0.867 \pm 0.018$ | $0.547 \pm 0.052$ |
| III | Path-only | Tuned | $0.957 \pm 0.024$ | $0.176 \pm 0.029$ | $0.867 \pm 0.018$ | $0.296 \pm 0.041$ |
| III | Full | Fixed | $0.532 \pm 0.081$ | $0.573 \pm 0.049$ | $0.903 \pm 0.015$ | $0.547 \pm 0.053$ |
| III | Full | Tuned | $0.857 \pm 0.032$ | $0.360 \pm 0.055$ | $0.903 \pm 0.015$ | $0.504 \pm 0.053$ |
| IV | Path-only | Fixed | $0.397 \pm 0.081$ | $0.597 \pm 0.100$ | $0.768 \pm 0.061$ | $0.469 \pm 0.058$ |
| IV | Path-only | Tuned | $0.963 \pm 0.028$ | $0.132 \pm 0.014$ | $0.768 \pm 0.061$ | $0.232 \pm 0.022$ |
| IV | Full | Fixed | $0.374 \pm 0.085$ | $\mathbf{0.632 \pm 0.088}$ | $0.804 \pm 0.054$ | $0.462 \pm 0.063$ |
| IV | Full | Tuned | $0.942 \pm 0.046$ | $0.134 \pm 0.015$ | $0.804 \pm 0.054$ | $0.234 \pm 0.023$ |

Note: AUC is threshold-independent; each AUC value applies to both the fixed- and tuned-threshold rows for the same context.

#### S2.3 Louvain-cluster-stratified prediction results

Louvain-cluster-stratified prediction results are reported in Supplementary Tables S2 and S3. The second-largest Louvain community produced the strongest fixed-threshold performance, particularly for the weighted path, whereas the largest Louvain community showed lower and more variable performance. As in the stage-stratified analysis, F1-based threshold tuning often increased recall but reduced precision and test-set F1-score relative to the fixed-threshold models, with the explanation likely being the same as above.

**Supplementary Table S2** Standard LSTM performance within the largest Louvain-derived community. Values are mean $\pm$ SD over 15 random seeds.

| Path | Context | Threshold | Recall | Precision | AUC | F1 |
|---|---|---|---|---|---|---|
| Unweighted | Path-only | Fixed | $0.294 \pm 0.078$ | $0.563 \pm 0.057$ | $0.755 \pm 0.028$ | $0.384 \pm 0.078$ |
| Unweighted | Path-only | Tuned | $0.903 \pm 0.074$ | $0.109 \pm 0.013$ | $0.755 \pm 0.028$ | $0.194 \pm 0.022$ |
| Unweighted | Full | Fixed | $0.288 \pm 0.079$ | $0.603 \pm 0.044$ | $0.785 \pm 0.019$ | $0.387 \pm 0.079$ |
| Unweighted | Full | Tuned | $0.909 \pm 0.090$ | $0.108 \pm 0.011$ | $0.785 \pm 0.019$ | $0.193 \pm 0.018$ |
| Weighted | Path-only | Fixed | $0.106 \pm 0.145$ | $0.272 \pm 0.273$ | $0.837 \pm 0.042$ | $0.135 \pm 0.156$ |
| Weighted | Path-only | Tuned | $0.944 \pm 0.090$ | $0.085 \pm 0.019$ | $0.837 \pm 0.042$ | $0.156 \pm 0.032$ |
| Weighted | Full | Fixed | $0.174 \pm 0.117$ | $0.488 \pm 0.374$ | $0.826 \pm 0.051$ | $0.238 \pm 0.153$ |
| Weighted | Full | Tuned | $0.849 \pm 0.189$ | $0.087 \pm 0.034$ | $0.826 \pm 0.051$ | $0.157 \pm 0.058$ |

Note: AUC is threshold-independent; each AUC value applies to both the fixed- and tuned-threshold rows for the same context.

**Supplementary Table S3** Standard LSTM performance within the largest Louvain-derived community. Values are mean ± SD over 15 random seeds.

| Path | Context | Threshold | Recall | Precision | AUC | F1 |
|---|---|---|---|---|---|---|
| Unweighted | Path-only | Fixed | 0.373 ± 0.037 | 0.587 ± 0.048 | 0.757 ± 0.021 | 0.455 ± 0.037 |
| Unweighted | Path-only | Tuned | 0.928 ± 0.044 | 0.139 ± 0.007 | 0.757 ± 0.021 | 0.242 ± 0.011 |
| Unweighted | Full | Fixed | 0.369 ± 0.032 | 0.626 ± 0.043 | 0.791 ± 0.018 | 0.464 ± 0.031 |
| Unweighted | Full | Tuned | 0.952 ± 0.027 | 0.136 ± 0.009 | 0.791 ± 0.018 | 0.238 ± 0.014 |
| Weighted | Path-only | Fixed | 0.599 ± 0.053 | 0.695 ± 0.075 | 0.860 ± 0.020 | 0.642 ± 0.056 |
| Weighted | Path-only | Tuned | 1.000 ± 0.000 | 0.158 ± 0.015 | 0.860 ± 0.020 | 0.273 ± 0.022 |
| Weighted | Full | Fixed | 0.593 ± 0.058 | 0.693 ± 0.077 | 0.875 ± 0.027 | 0.638 ± 0.059 |
| Weighted | Full | Tuned | 0.959 ± 0.044 | 0.161 ± 0.020 | 0.875 ± 0.027 | 0.275 ± 0.030 |

Note: AUC is threshold-independent; each AUC value applies to both the fixed- and tuned-threshold rows for the same context.
